# Trait level somatic arousal modulates fMRI neural synchrony to naturalistic stimuli

**DOI:** 10.1101/2023.09.27.559823

**Authors:** K. Klamer, J. Craig, K. Sullivan, C. Haines, C. Ekstrand

## Abstract

Somatic arousal refers to the physiological and bodily responses that occur in reaction to different emotional and psychological stimuli and is a crucial component of the fight or flight response. Symptoms associated with higher levels of somatic arousal such as higher heart and respiration rates have been shown to impact the blood oxygenation level dependent (BOLD) signal during functional magnetic resonance imaging (fMRI) studies. Differences in baseline levels of somatic arousal may therefore modulate the brain’s response to incoming stimuli during fMRI. Previous studies typically investigate somatic arousal as a state, rather than as a trait, in which some individuals are more likely to have heightened physiological responses to psychological stimuli, causing the neurological mechanisms behind baseline somatic arousal levels to remain poorly understood. The current study seeks to identify how differing levels of baseline somatic arousal modulate neural synchrony in response to an audiovisual film. We hypothesize that individuals with higher levels of somatic arousal will show overall heightened neural synchrony in response to a complex audiovisual stimulus. We identified that higher levels of somatic arousal are associated with widespread neural synchrony across the brain, including frontal gyri, parietal and temporo-occipital cortices. Taken together, this research suggests that baseline somatic arousal levels should be measured during naturalistic fMRI paradigms, as baseline somatic arousal levels may have a profound influence on synchronous neural activity.

---

Somatic arousal refers to the physiological and bodily responses that occur in reaction to emotional and psychological stimuli. It is a fundamental aspect of the “fight or flight” response, which is the body’s natural reaction to perceived threats or stressors (Salsman et al., 2013). It can be triggered by external stimuli with negative emotional valence, such as fear, stress, and anxiety (Steimer, 2022), as well as positive emotional valence, including excitement and joy (Vergallito et al., 2019). Physical manifestations of somatic arousal include increased heart rate, blood pressure, glandular secretion, body temperature, muscle twitches/trembling, nausea, and shortness of breath (McCorry, 2007; Passatore & Roatta, 2006; Pappas, 2003; Stockhorst et al., 1993; Russo et al., 2017). Somatic arousal is a critical mechanism that helps humans respond quickly to potential threats or challenges. However, chronic or excessive somatic arousal has been linked with negative mental health outcomes, including anxiety, post-traumatic stress disorder, neuroticism, and emotional dysregulation (Shepherd & Wild, 2014) through its association with intensified emotions of fear (Storbeck & Clore, 2008) and misinterpretation of ambiguous situations (Ohst & Tuschen-Caffier, 2020). Physical health outcomes associated with heightened levels of somatic arousal include cardiovascular disease and immune system suppression (Dienstibier, 1989). Thus, understanding somatic arousal and its effects can be crucial in managing stress, anxiety, and other emotional responses.

Acute changes in somatic arousal levels can also influence cognition and shape how people respond to everyday tasks. For example, increased levels of somatic arousal increase attentional selectivity (Sanbonmatsu & Kardes, 1998) and predict time distortion in the context of highly arousing negative stimuli, where increases in physiological arousal lengthen the perceived duration of events (Ogden et al., 2019). Somatic arousal has further been shown to impact memory retrieval and working memory. Buchanan et al. (2006) investigated the influence of autonomic arousal and memory for emotional words and found that the words that elicited greater autonomic activity were better remembered than the words that did not, suggesting that acute somatic arousal may produce improvements in memory retrieval. Somatic arousal has been hypothesized to play a crucial role in the visceral feedback of emotion, where increased somatic arousal enhances emotions such as fear, anger, and love, and decreased somatic arousal can reduce the intensity of emotional experiences (Laird, 2007). High empathy scores have been linked to increased autonomic arousal at the onset of emotional stimuli (Bogdanov et al., 2013), and feelings of familiarity may also stem from autonomic arousal associated with cognitive resource allocation (Morris et al., 2008). Therefore, the impact of acute changes in somatic arousal on cognition is far-reaching and may influence both top-up and bottom-down neural processing.

The brain regions generally associated with somatic arousal are the brain regions associated with state-anxiety, which is associated with excessive somatic arousal. Connections between the postcentral gyrus and left cerebellum are positively correlated with state anxiety, whereas connections between the postcentral gyrus, the left inferior frontal cortex and left medial superior frontal cortex are negatively associated with state anxiety (Li et al., 2019). Another study investigating arousal as a function of wakefulness has suggested that regions involved in sensory processing, including the thalamus, brainstem, postcentral gyrus, and visual/auditory cortices are associated with higher arousal levels (McGinley, 2020). While these studies are not specific to heightened somatic arousal, they provide a valuable starting point for understanding somatic arousal in the brain.

To date, little research has focused on the neural mechanisms underlying differing levels of baseline somatic arousal. Most of the literature surrounding somatic arousal focuses on acute symptoms (Satpute et al., 2019; Fan et al., 2015), rather than as a trait in which some individuals are more likely to have more pronounced somatic arousal responses. Furthermore, studies investigating somatic arousal tend to use sparse tasks, such as viewing emotional photos (Hillman et al., 2004) or completing memory tasks (Sawai et al., 2015), and behavioural measures of somatic arousal, such as measuring heart rate variability (Graham et al., 2000). However, simple, sparse tasks, such as photo viewing and memory tasks, do not reflect the complexity of stimuli encountered in real-life. These studies typically isolate specific components of multimodal integration through simplified stimuli (i.e., visual input, emotional processes, stress-inducing auditory input), although multimodal integration is most likely essential for coherent perceptual experience (Sonsukare et al., 2019).

Naturalistic paradigms, which use stimuli such as dynamic videos, speech, and music, have been increasingly employed in neuroimaging studies in recent years to study more true-to-life sensory experience. These paradigms employ the dynamic, rich and multimodal stimuli that better represent our lived experience. Naturalistic paradigms, while still delivered in a laboratory setting, provide an approximation to how we encounter stimuli in everyday life (Sonkusare et al., 2019). Using naturalistic paradigms provides analytic flexibility due to the stimuli requiring continuous, real-time integration of dynamic streams of information (Bottenhorn et al., 2018). Naturalistic paradigms do not necessarily mimic the natural world; rather, they aim to evoke more naturalistic patterns of neural responses (Vanderwal et al., 2019). For large, overarching topics such as somatic arousal, using naturalistic stimuli may be a more appropriate measure to provide a well-rounded picture of how somatic arousal influences everyday life.

Intersubject correlation analysis (ISC) is a technique often used to analyze complex, naturalistic fMRI data. ISC examines correlations in hemodynamic responses across the time course of an audiovisual stimulus to identify neural synchrony, i.e., the neural activity that is shared between subjects (Pajula et al., 2012). Due to the influences from blood gas levels and blood pressure on tissue oxygen demand and blood vessel volume, changes in blood pressure or oxygenation can lead to changes in fMRI signals (Macey et al., 2016). As heightened somatic arousal is associated with increases in heart rate and respiration rate (McCorry, 2007), different levels of baseline somatic arousal may modulate changes in neural synchrony. This may further be associated with the changes in cognition correlated with somatic arousal discussed previously. However, it is currently unknown how trait somatic arousal modulates neural synchrony.

The current study seeks to investigate how differing levels of trait somatic arousal impact neural synchrony to naturalistic stimuli. Specifically, we seek to examine differences in neural synchrony between a group of participants with relative low self-reported baseline somatic arousal and high baseline somatic arousal in response to a full-length audiovisual movie. To do so, we used preprocessed fMRI data from the Naturalistic Neuroimaging Database (NNDb) v2.0 (Aliko et al., 2020) and separated participants based on their somatic arousal scores, as quantified using the NIH Toolbox 2.0 Fear-Somatic Arousal instrument (Salsman et al., 2013). As somatic arousal leads to physiological changes that are associated with several cognitive traits, we hypothesize that individuals with higher levels of somatic arousal will show overall heightened neural synchrony in response to a complex audiovisual stimulus.

## Methods

### Participants and fMRI data

We used fMRI data from the publicly available Naturalistic Neuroimaging Database (NNDb, v2.0; Aliko et al., 2020). We selected 20 participants (10 females/10 males, aged 19-53, mean age of 27.7 years) who watched the feature-length audiovisual film *500 Days of Summer* (Webb, 2009; duration ∼ 95 minutes) during fMRI. All participants were right-handed, native English speakers, with no history of neurological/psychiatric illnesses, no hearing impairments, unimpaired or corrected vision, and did not take medication. Full fMRI preprocessing details can be found in Aliko et al. (2020). Briefly, all data was acquired on a 1.5T Siemens MAGNETOM Avanto, and functional data was acquired using a multiband echo-planar imaging (EPI) sequence (TR: 1s, TE: 58.4ms). We used version 2.0 of the NNDb (Aliko et al., 2020), which, in comparison to v1.0, has improved normalization and standardization of the data, resulting in ‘cleaner’ preprocessed data and more robust statistics. Aliko et al., (2020) used the *afni_proc.py* pipeline from AFNI (Cox & Hyde, 1997; Cox, 1999) for preprocessing, correcting for slice-timing differences, despiking, correcting for motion, spatially aligning the data to an MNI template with a resampling size of 3×3×3mm^3^, and correcting for timing to align the fMRI time series and the film. The authors of the NNDb (v2.0) (Aliko et al., 2020) obtained approval by the ethics committee of University College London and participants provided written informed consent to take part in the study and share their anonymized data.

### Behavioural Questionnaires

Following the MRI imaging session, participants completed the majority of the National Institute of Health (NIH) Toolbox, which validates measures of sensory, motor, cognitive and emotional processing to measure individual differences (Gershon et al., 2013). In this study, we used the Fear-somatic arousal (FSA) scores quantified using the NIH Toolbox 2.0 Fear-Somatic Arousal (18+; Gershon et al., 2013) questionnaire. We used a median split analysis to separate participants into low FSA and high FSA groups based on their provided T-score, resulting in 10 participants in each group. We used SPSS (version 27; IBM Corp., 2020) to run an independent sample t-test between participants in each FSA group (i.e., high-high and low-low FSA) and age.

### Intersubject Correlation Analyses

Intersubject correlation analyses (ISC) were run for all unique pairs of participants, which produced 190 (*n*\*(*n*-1)/2, where n = 20) unique ISC maps. ISC is a model-free approach used to analyze complex fMRI data acquired in naturalistic, audiovisual stimulus environments (Kauppi et al., 2010). It allows us to measure shared content across experimental conditions by filtering out subject-specific signals and revealing voxels with a consistent, stimulus-evoked response time series across subjects (Nastase et al., 2019). It does this by calculating pairwise correlation coefficients between all pairs of participants for each voxel throughout the brain. We separated the resulting 190 ISC maps into three groups based on FSA grouping (i.e., low-low FSA pairwise correlations grouped, high-high pairwise correlations grouped, low-high pairwise correlations grouped). This resulted in 45 (*n*\*(*n*-1)/2, where n = 10) unique ISC maps for both the high (i.e., high-high) and low (i.e., low-low) FSA groups, and 100 between group pairs (i.e., low-high).

### FMRI Analysis

After running ISCs on all pairs of participants, we used the Linear Mixed Effects Model (LME) implemented via the *3dISC* module in AFNI (Cox & Hyde, 1997; Cox, 1999; as described by Chen et al., 2020) to obtain the following contrasts: low-low vs. low-high FSA, low-low vs. high-high FSA, and high-high vs. low-high FSA. LME is a parametric method based on the general linear model (GLM) and is used to model the relationship between the fMRI time series data and the experimental conditions at each voxel, accounting for the complex covariance structure of ISC data. As LME accounts for variability in the data due to random effects, it allows for more precise estimation of fixed effects (Koerner & Zhang, 2017). We modeled the following contrasts: ISC in the low FSA group (low-low FSA), ISC in the high FSA group (high-high FSA), ISC between the high and low groups (low-high FSA), comparison of ISC between the low and high groups (low-low vs. high-high FSA), comparison of ISC between the low group and between group ISC (low-low vs. low-high), and comparison of ISC between the high group and between group ISC (high-high vs. low-high). Significant ISCs for the group average contrasts (i.e.., high-high FSA and low-low FSA), were defined using a voxelwise false discovery rate (FDR) of *q* < 0.0001. Significant ISCs for the group comparisons contrasts (i.e., low-low FSA vs. high-high FSA; low-low FSA vs. low-high FSA; high-high vs. low-high FSA) were defined using a voxelwise FDR of *q* < 0.05. Significant results were transformed into surface space for visualization purposes only.

## Results

### Behavioral Questionnaires

We used SPSS (version 27; IBM Corp., 2020) to run an independent sample t-test between participants in the high and low FSA groups to determine whether age differed between the two groups. No significant differences were found *t*(18) = .922, *p* = .369.

### Group Averages

Results from the low-low FSA and high-high FSA group average contrasts showed widespread ISCs across the occipital, temporal, and parietal cortices, which is in line with previous research (Güçlütürk et al., 2018). Full results for low-low FSA and high-high FSA group averages can be found in Supplementary Materials A.

### Low-low vs. High-high FSA contrasts

#### Low-low > High-high FSA

Results from the low-low FSA > high-high FSA are shown in Figure 1 and Supplementary Materials B. The Low-low FSA group showed significantly greater synchrony than the High-high FSA group in the left pons of the brainstem, bilateral superior temporal gyrus, bilateral superior parietal lobule, bilateral lateral occipital cortex, and bilateral occipital pole.

**Figure 1.**
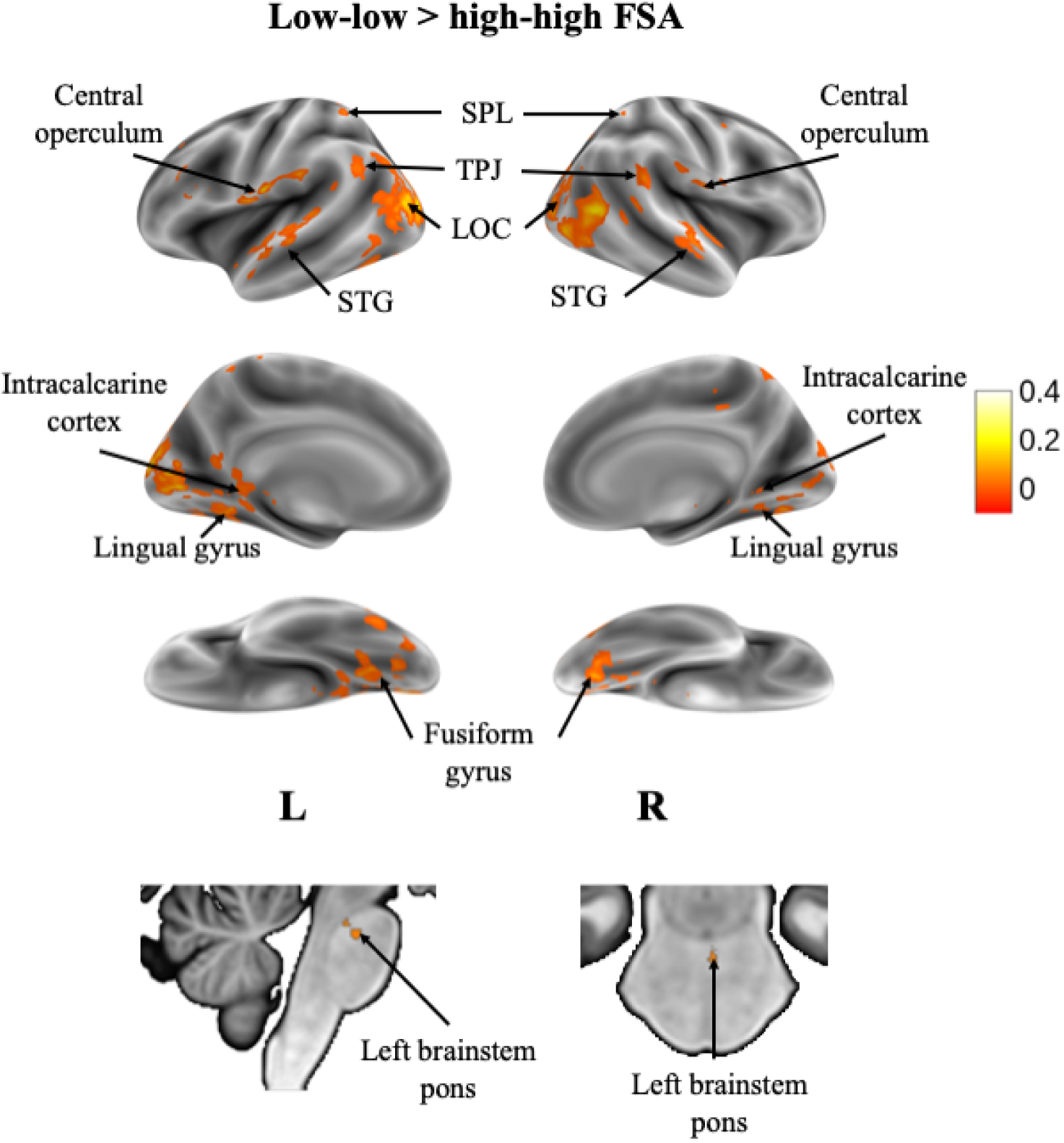
Voxels showing significant ISC across the time course of the audiovisual stimulus in participants within the low-low FSA group (*n* = 10) in comparison to participants within the high-high FSA group (*n* = 10). Results are displayed as a voxelwise false-discovery rate (FDR) threshold of *q* = 0.05.

#### High-high > Low-low FSA

Results from the High-high > Low-low FSA contrast are shown in Figure 2 and Supplementary Materials B. Overall, the high-high FSA group showed significantly greater synchrony than the low-low FSA group across a wide array of the cortex, including the bilateral orbitofrontal cortex (OFC), bilateral precuneus, bilateral posterior cingulate cortex (PCC), bilateral inferior and middle frontal gyri, bilateral middle and superior temporal gyri, bilateral occipital pole, bilateral planum temporale, bilateral pre/postcentral gyri, and bilateral cerebellar cortices.

**Figure 2.**
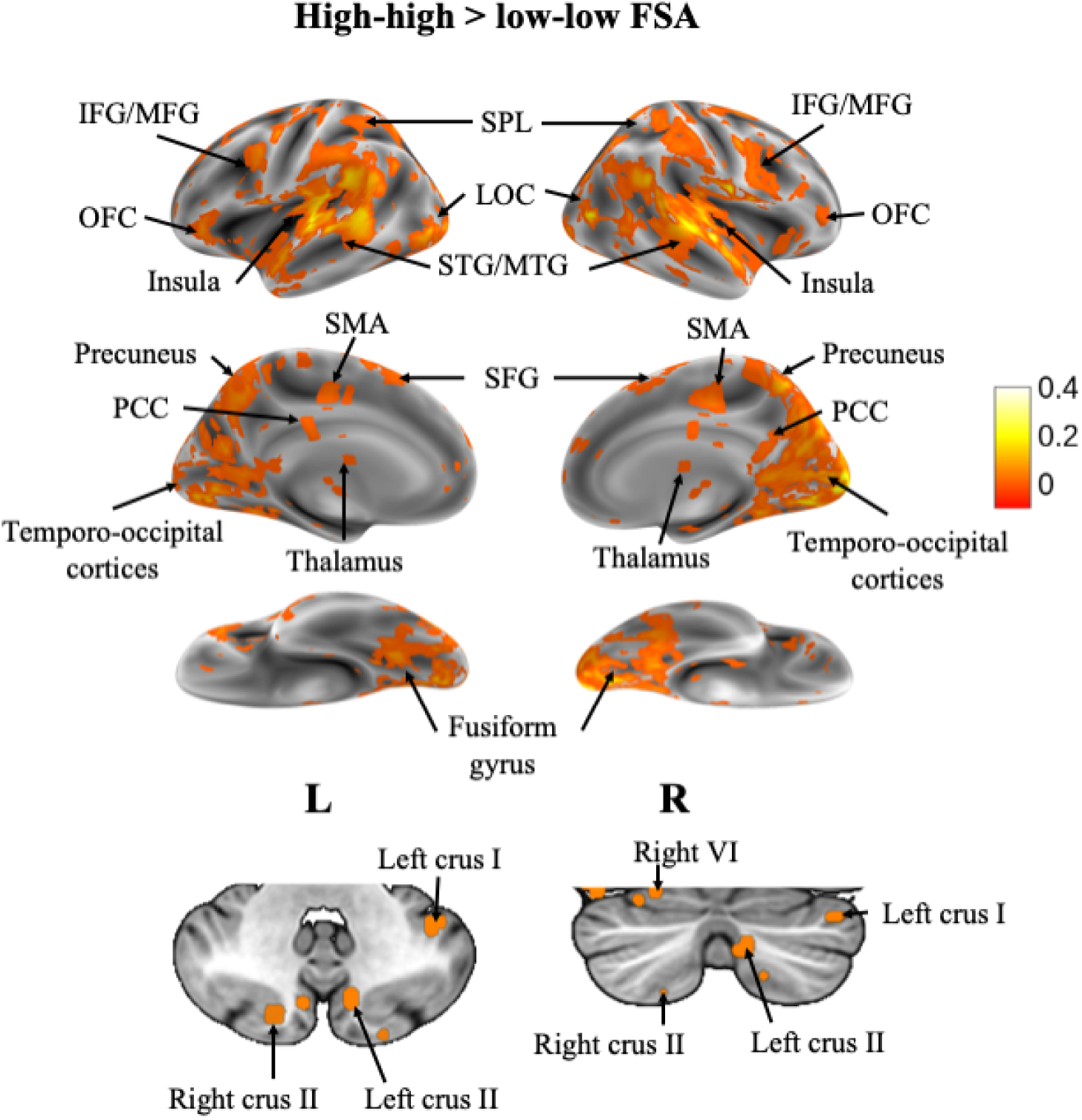
Voxels showing significant ISC across the time course of the audiovisual stimulus in participants within the high-high FSA group (*n* = 10) in comparison to participants within the low-low FSA group (*n* = 10). Results are displayed as a voxelwise false-discovery rate (FDR) threshold of *q* = 0.05.

### Comparing low and high group ISC with between group ISC

#### Low-low > Low-high FSA

Results from the Low-low > Low-high FSA contrast are shown in Figure 3 and Supplementary Materials C. This contrast showed limited neural synchrony, with one voxel showing heightened neural synchrony in the left planum temporale, and one voxel showing heightened neural synchrony in the left occipital pole.

**Figure 3.**
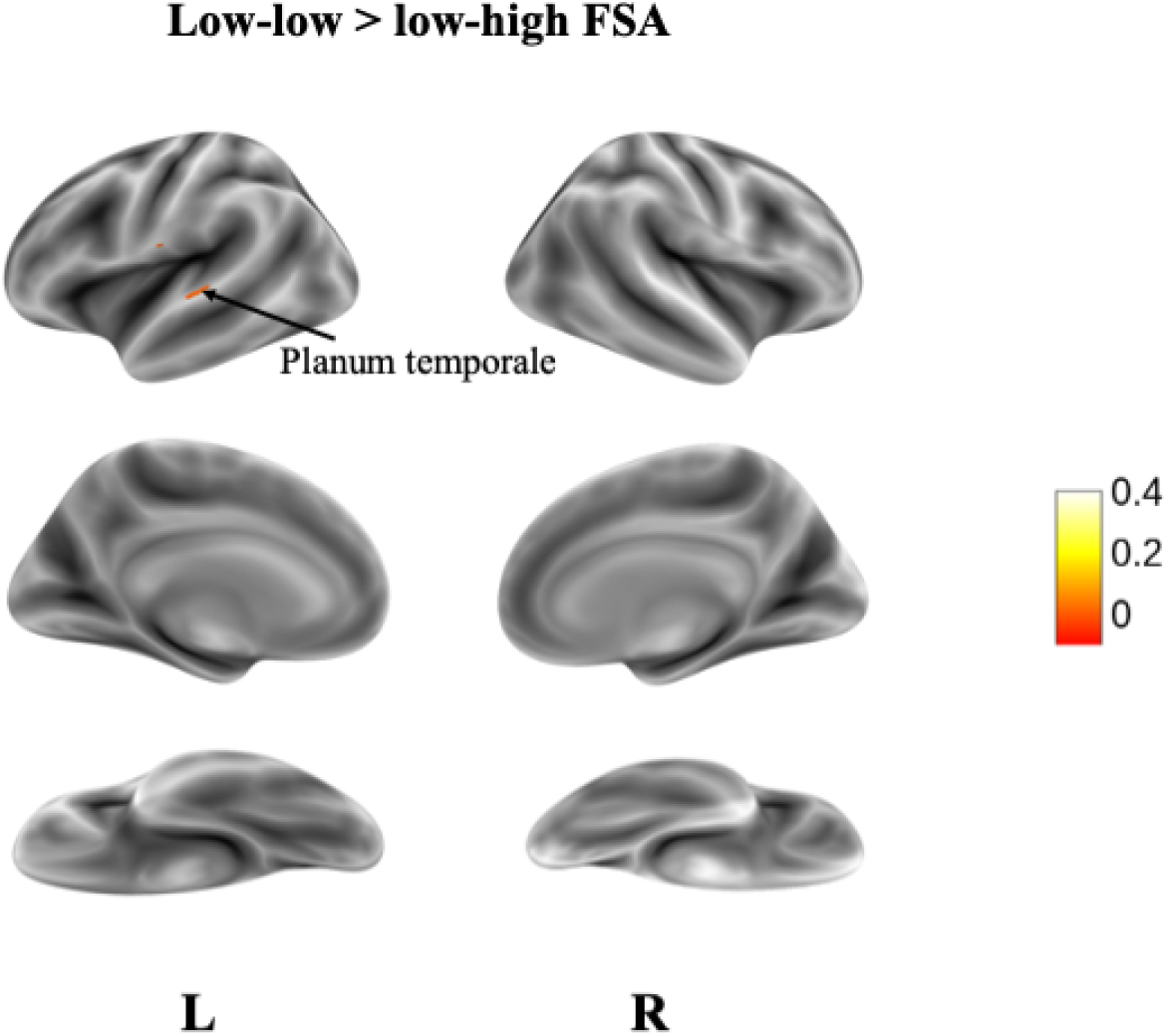
Voxels showing significant ISC across the time course of the audiovisual stimulus in participants within the low-low FSA group in comparison to participants within the low-high FSA group. Results are displayed as a voxelwise false-discovery rate (FDR) threshold of *q* = 0.05.

#### High-high > Low-high FSA

Results from the High-high > Low-high FSA contrast are shown in Figure 4 and Supplementary Materials C. Overall, the high-high FSA group showed significantly greater synchrony than the low-high FSA group across a wide array of the cortex, including the bilateral orbitofrontal cortex (OFC), bilateral precuneus, bilateral temporal parietal junction (TPJ), bilateral middle and superior temporal gyri, bilateral insula, bilateral superior parietal lobule, bilateral occipital pole, bilateral planum temporale, bilateral precentral gyri, and bilateral cerebellar cortices.

**Figure 4.**
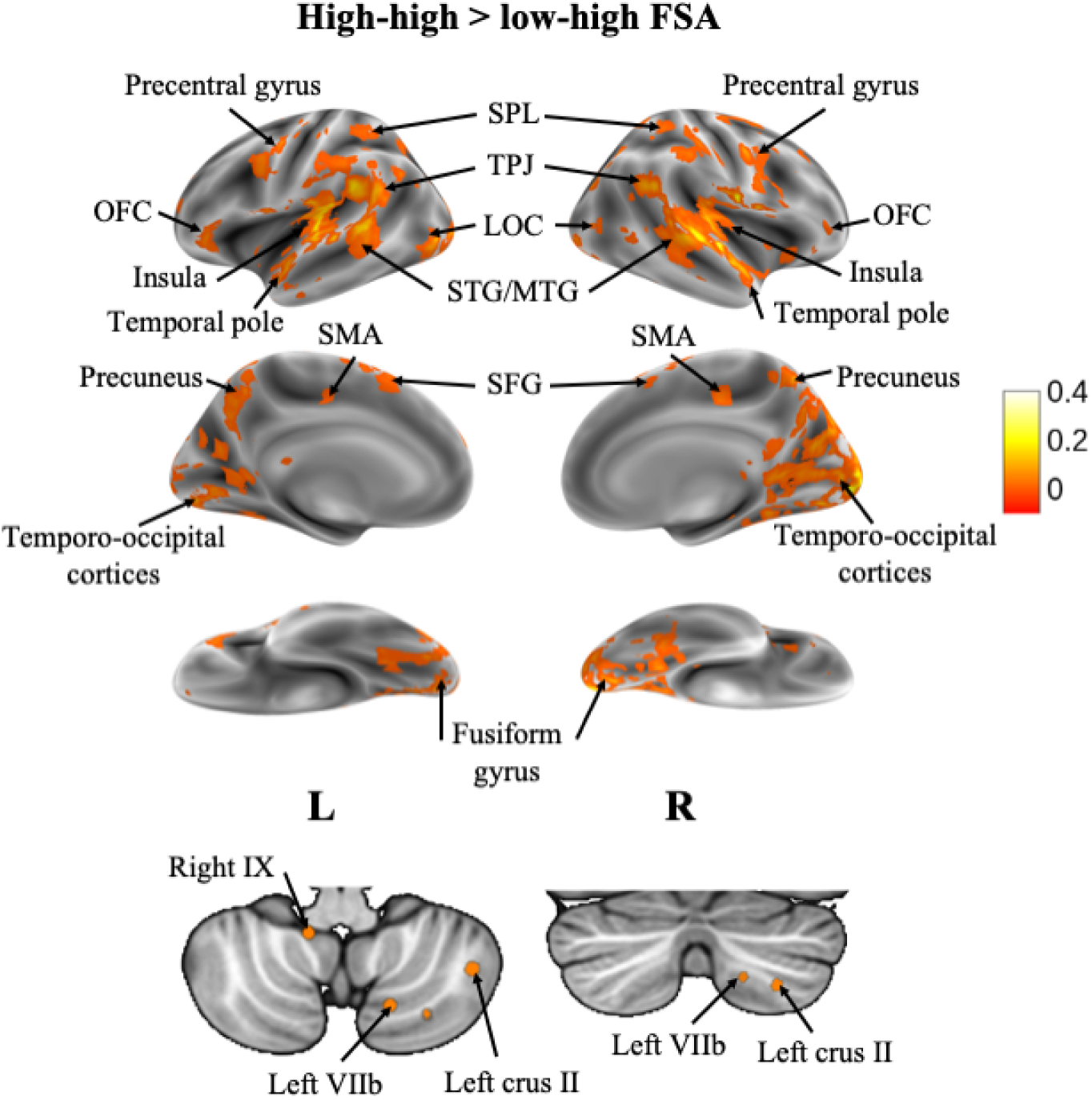
Voxels showing significant ISC across the time course of the audiovisual stimulus in participants within the high-high FSA group in comparison to participants within the low-high FSA group. Results are displayed as a voxelwise false-discovery rate (FDR) threshold of *q* = 0.05.

## Discussion

This study sought to examine differences in neural synchrony to a complex audiovisual stimulus based on levels of baseline somatic arousal. Neural activity has been shown to be sensitive to the nature of the stimulus, and results from this study have confirmed that trait-level individual differences in baseline somatic arousal result in substantial variations in neural synchrony in response to a naturalistic stimulus. Consistent with our hypothesis, in comparison to individuals with lower levels of somatic arousal, higher levels of somatic arousal were associated with widespread neural synchrony across wide expanses of the cortex, including regions associated with emotional processing, the dorsal attention network, and working memory.

The regions that showed the most pronounced difference, with higher synchronization in the high-high FSA group, include visual and auditory processing areas, such as the lateral occipital cortex, superior temporal gyrus, and medial temporo-occipital cortices. This indicates that individuals with higher levels of somatic arousal processed the visual and auditory components of the film more similarly than participants in the low-low and low-high FSA groups. This effect could be due to the increases in sustained attention that are associated with higher levels of somatic arousal, as studies have shown that attention modulates the strength and speed of audiovisual processing (Seijdel et al., 2023), or it could be a result of the proposed heightened brain-wide excitability associated with higher levels of somatic arousal (Raut et al., 2021). More research is needed to fully elucidate changes in visual and auditory sensory processing associated with heightened somatic arousal levels.

Other regions that were found to have higher synchronization in the high-high FSA group compared to the low-low and low-high FSA groups include the temporal parietal junction, frontal eye fields, superior parietal lobule, intraparietal sulcus, and ventral premotor cortex, which comprise the dorsal attention network (DAN; Spreng et al., 2017). The DAN is a bilateral network that focuses human attention (Farrany & Uddin, 2015), and sustained attention is modulated by physiological states of arousal (Pinggal et al., 2022). Further, higher levels of attention have been shown to enhance neural synchrony during narrative processing (Jääskeläinen et al., 2020). Therefore, the results of the present study emphasize the findings that sustained attention is influenced by somatic arousal levels, and suggest that this influence may be modulated by a differential BOLD response throughout key regions of the DAN.

Brain regions involved in working memory were found to have higher synchronization in the high-high FSA group compared to the low-low FSA group, including the dorsolateral prefrontal cortex (dlPFC), ACC, and the parietal cortex (Chai et al., 2018). This indicates that the working memory network behaved more similarly during movie-watching in participants with higher levels of somatic arousal. Somatic arousal has been found to be correlated with working memory outcomes, with emotional arousal enhancing memory for central details of an event (Hulse et al., 2007) and the slower forgetting of arousing words (Sharot et al., 2004). Further, it has been suggested that arousal may act as a cue for arousal-related material in memory (Clark et al., 1983), overall indicating that somatic arousal can play a role in memory retention and the encoding of emotional and arousing information. Higher levels of working memory have further been associated with neural synchrony, with higher working memory being associated with increased synchrony in brain regions such as the vlPFC and frontoparietal network (Saliasi et al., 2014), regions observed in the current study. Further, an EEG study found that working memory may be partly subserved by synchronization in a bilateral frontal network (Serrien et al., 2004). Consequently, the current study provides support for increases in working memory associated with higher levels of somatic arousal in response to an audiovisual film, as demonstrated by increased neural synchrony in regions associated with working memory.

FMRI is sensitive to changes in vascular resistance and blood pressure, as it relies on a localized cerebral blood flow response to changes in cortical neuronal activity (Duyn et al., 2020). As individuals with higher baseline somatic arousal are more likely to experience heightened states of physiological arousal in response to an audiovisual film, it is possible that the associated increase in blood flow and respiration rate produced similar changes in neuronal activity across participants in the high-high FSA group, resulting in an increase in neural synchrony. Further, Raut et al. (2021) used fMRI to demonstrate that fluctuations in arousal level are associated with slowly propagating parallel waves of activity throughout the neocortex, thalamus, striatum, and cerebellum, and demonstrated that features of spontaneous fMRI signal fluctuations can be parsimoniously accounted for by these waves. They further suggest that traveling waves spatiotemporally pattern brain-wide excitability in relation to arousal. If we extend this finding to our results, this could indicate that higher levels of baseline somatic arousal are indicative of heightened brain-wide excitability, as demonstrated by increased neural synchrony throughout the brain.

Several cognitive traits are associated with higher levels of somatic arousal, one being the more intense experience of emotions. Not only does the experience of emotions enhance the synchrony of brain activity across individuals (Nummenmaa et al., 2012), but highly arousing emotions are associated with increased BOLD signal intensities in regions such as the left thalamus, bilateral parahippocampal gyrus, and bilateral premotor cortex (Colibazzi et al., 2010). As we observed heightened neural synchrony across the brain and specifically in these regions associated with highly arousing emotions within the high-high FSA group, it is plausible that this increased physiological response during the film as a result of higher baseline somatic arousal levels may have contributed to a corresponding BOLD response, potentially manifesting as the enhanced intensity of emotional experiences.

This study aimed at uncovering differential neural synchrony associated with varying levels of baseline somatic arousal levels during movie watching, however, there are several limitations to be discussed. Due to the use of an online database, we were unable to have control over the stimuli that were presented to the participants, or the measures of baseline somatic arousal implemented. This introduces a limitation as somatic arousal was assessed solely using self-report measures, rather than combining self-report measures and reliable physiological indices. Additionally, the NIH Toolbox 2.0 Fear-Somatic Arousal questionnaire only asks about somatic arousal symptoms experienced over the past seven days, which allows the possibility for subjective life circumstances to impact the self-reported baseline somatic arousal assessment. Additionally, the data used in these experiments was collected using a 1.5T MRI, which has a lower signal-to-noise ratio and a lower spatial specificity than MRIs with a stronger field strength. Future research should aim at combining self-report and reliable physiological measures to assess somatic arousal and should be investigated using higher field strengths to acquire such data.

In conclusion, this study demonstrated that baseline somatic arousal may modulate widespread neural synchrony during movie-watching. We identified neural correlates associated with sustained attention, emotional processing, and working memory that may be associated with higher levels of somatic arousal. We further provided evidence that can be used to support the notion that higher somatic arousal is associated with brain-wide excitability. The findings collectively emphasize that during naturalistic fMRI paradigms, baseline somatic arousal levels should be measured and controlled for as this trait may play a crucial role in modulating neural synchrony in response to an audiovisual film.

## Author Contributions

The primary author of this study is K.K. The co-authors of this study are J. C., K. S., C. H., and C.E.

## Data availability statement

The preprocessed fMRI data used in this study is openly available and can be downloaded from the Naturalistic Neuroimaging Database (version 2.0.0; https://openneuro.org/datasets/ds002837/versions/2.0.0).

## References

Aly, M., Chen, J., Turk-Browne, N. B., & Hasson, U. (2018). Learning naturalistic temporal structure in the posterior medial network. Journal of Cognitive Neuroscience, 30(9), 1345–1365.

Bogdanov, V. B., Bogdanova, O. V., Gorlov, D. S., Gorgo, Y. P., Dirckx, J. J., Makarchuk, M. Y.,… & Critchley, H. (2013). Alexithymia and empathy predict changes in autonomic arousal during affective stimulation. Cognitive and Behavioral Neurology, 26(3), 121–132.

Bottenhorn, K. L., Flannery, J. S., Boeving, E. R., Riedel, M. C., Eickhoff, S. B., Sutherland, M. T., & Laird, A. R. (2018). Cooperating yet distinct brain networks engaged during naturalistic paradigms: A meta-analysis of functional MRI results. Network Neuroscience, 3(1), 27–48.

Broderick, J. E., Gold, M. S., Amin, M. M., & Gold, A. R. (2014). The association of somatic arousal with the symptoms of upper airway resistance syndrome. Sleep Medicine, 15(4), 436–443.

Buchanan, T. W., Etzel, J. A., Adolphs, R., & Tranel, D. (2006). The influence of autonomic arousal and semantic relatedness on memory for emotional words. International journal of psychophysiology, 61(1), 26–33.

Chai, W. J., Abd Hamid, A. I., & Abdullah, J. M. (2018). Working memory from the psychological and neurosciences perspectives: a review. Frontiers in psychology, 9, 401.

Chen, Gang, Paul A. Taylor, Yong Wook Shin, Richard C. Reynolds, and Robert W. Cox. 2017. “Untangling the Relatedness among Correlations, Part II: Inter-Subject Correlation Group Analysis through Linear Mixed-Effects Modeling.” NeuroImage 147 (February): 825–40.

Chen, G., Taylor, P. A., Qu, X., Molfese, P. J., Bandettini, P. A., Cox, R. W., & Finn, E. S. (2020). Untangling the relatedness among correlations, part III: inter-subject correlation analysis through Bayesian multilevel modeling for naturalistic scanning. NeuroImage, 216, 116474.

Clark, M. S., Milberg, S., & Ross, J. (1983). Arousal cues arousal-related material in memory: Implications for understanding effects of mood on memory. Journal of Verbal Learning and Verbal Behavior, 22(6), 633–649.

Cox, R. W. (1996). AFNI: software for analysis and visualization of functional magnetic resonance neuroimages. Computers and Biomedical research, 29(3), 162–173.

Cox, R. W., & Hyde, J. S. (1997). Software tools for analysis and visualization of fMRI data. NMR in Biomedicine: An International Journal Devoted to the Development and Application of Magnetic Resonance In Vivo, 10(4LJ5), 171–178.

Critchley, H. D. (2002). Electrodermal responses: what happens in the brain. The Neuroscientist, 8(2), 132–142.

Critchley, H. D., Rotshtein, P., Nagai, Y., O’Doherty, J., Mathias, C. J., & Dolan, R. J. (2005). Activity in the human brain predicting differential heart rate responses to emotional facial expressions. Neuroimage, 24(3), 751–762.

Diek, D., Smidt, M. P., & Mesman, S. (2022). Molecular organization and patterning of the medulla oblongata in health and disease. International Journal of Molecular Sciences, 23(16), 9260.

Dienstbier, R. A. (1989). Arousal and physiological toughness: implications for mental and physical health. Psychological review, 96(1), 84.

Duyn, J. H., Ozbay, P. S., Chang, C., & Picchioni, D. (2020). Physiological changes in sleep that affect fMRI inference. Current Opinion in Behavioral Sciences, 33, 42–50.

Fan, J., Xu, P., Van Dam, N. T., Eilam-Stock, T., Gu, X., Luo, Y. J., & Hof, P. R. (2012). Spontaneous brain activity relates to autonomic arousal. Journal of neuroscience, 32(33), 11176–11186.

Farrant, K., & Uddin, L. Q. (2015). Asymmetric development of dorsal and ventral attention networks in the human brain. Developmental cognitive neuroscience, 12, 165–174.

Finn, E. S., Corlett, P. R., Chen, G., Bandettini, P. A., & Constable, R. T. (2018). Trait paranoia shapes inter-subject synchrony in brain activity during an ambiguous social narrative. Nature communications, 9(1), 2043.

Fish, D. R., Gloor, P., Quesney, F. L., & Oliver, A. (1993). Clinical responses to electrical brain stimulation of the temporal and frontal lobes in patients with epilepsy: pathophysiological implications. Brain, 116(2), 397–414.

Graham, C., Sastre, A., Cook, M. R., & Kavet, R. (2000). Heart rate variability and physiological arousal in men exposed to 60 Hz magnetic fields. Bioelectromagnetics: Journal of the Bioelectromagnetics Society, The Society for Physical Regulation in Biology and Medicine, The European Bioelectromagnetics Association, 21(6), 480–482.

Güçlütürk, Y., Güçlü, U., van Gerven, M., & van Lier, R. (2018). Representations of naturalistic stimulus complexity in early and associative visual and auditory cortices. Scientific reports, 8(1), 3439.

Hillman, C. H., Rosengren, K. S., & Smith, D. P. (2004). Emotion and motivated behavior: postural adjustments to affective picture viewing. Biological psychology, 66(1), 51–62.

Hulse, L. M., Allan, K., Memon, A., & Read, J. D. (2007). Emotional arousal and memory: A test of the poststimulus processing hypothesis. The American journal of psychology, 120(1), 73–90.

IBM Corp. Released 2020. IBM SPSS Statistics for Windows, Version 27.0. Armonk, NY: IBM Corp

Leech, R., & Sharp, D. J. (2014). The role of the posterior cingulate cortex in cognition and disease. Brain, 137(1), 12–32

Kauppi, J. P., Jääskeläinen, I. P., Sams, M., & Tohka, J. (2010). Inter-subject correlation of brain hemodynamic responses during watching a movie: localization in space and frequency. Frontiers in neuroinformatics, 5.

Koerner, Tess K., and Yang Zhang. 2017. “Application of Linear Mixed-Effects Models in Human Neuroscience Research: A Comparison with Pearson Correlation in Two Auditory Electrophysiology Studies.” Brain Sciences 2017, Vol. 7, Page 26 7 (3): 26.

Laird, J. D. (2007). Feelings: The perception of self. Oxford University Press

Levenson, R. W. (2014). The autonomic nervous system and emotion. Emotion review, 6(2), 100–112.

Li, X., Zhang, M., Li, K., Zou, F., Wang, Y., Wu, X., & Zhang, H. (2019). The altered somatic brain network in state anxiety. Frontiers in psychiatry, 10, 465.

Macey, P. M., Ogren, J. A., Kumar, R., & Harper, R. M. (2016). Functional imaging of autonomic regulation: methods and key findings. Frontiers in neuroscience, 9, 513.

Marko, M., & Riečanský, I. (2018). Sympathetic arousal, but not disturbed executive functioning, mediates the impairment of cognitive flexibility under stress. Cognition, 174, 94–102.

McCorry, L. K. (2007). Physiology of the autonomic nervous system. American journal of pharmaceutical education, 71(4).

McGinley, M. J. (2020). Brain states: Sensory modulations all the way down. Current Biology, 30(20), R1263–R1266.

Morris, A. L., Cleary, A. M., & Still, M. L. (2008). The role of autonomic arousal in feelings of familiarity. Consciousness and cognition, 17(4), 1378–1385.

Nastase, S. A., Gazzola, V., Hasson, U., & Keysers, C. (2019). Measuring shared responses across subjects using intersubject correlation. Social Cognitive and Affective Neuroscience, 14(6), 667–685.

Nummenmaa, L., Glerean, E., Viinikainen, M., Jääskeläinen, I. P., Hari, R., & Sams, M. (2012). Emotions promote social interaction by synchronizing brain activity across individuals. Proceedings of the National Academy of Sciences, 109(24), 9599–9604.

Ogden, R. S., Henderson, J., McGlone, F., & Richter, M. (2019). Time distortion under threat: Sympathetic arousal predicts time distortion only in the context of negative, highly arousing stimuli. PloS one, 14(5), e0216704.

Oppenheimer, S. M., Gelb, A., Girvin, J. P., & Hachinski, V. C. (1992). Cardiovascular effects of human insular cortex stimulation. Neurology, 42(9), 1727–1727.

Pajula, J., Kauppi, J.-P., & Tohka, J. (2012). Inter-Subject Correlation in fMRI: Method Validation against Stimulus-Model Based Analysis. PLoS ONE, 8(8), 41196. 10.1371/journal.pone.0041196.

Pappas Jr, D. G. (2003). Autonomic related vertigo. The Laryngoscope, 113(10), 1658–1671.

Pinggal, E., Dockree, P. M., O’Connell, R. G., Bellgrove, M. A., & Andrillon, T. (2022). Pharmacological manipulations of physiological arousal and sleep-like slow waves modulate sustained attention. Journal of Neuroscience, 42(43), 8113–8124.

Pollatos, O., Schandry, R., Auer, D. P., & Kaufmann, C. (2007). Brain structures mediating cardiovascular arousal and interoceptive awareness. Brain research, 1141, 178–187.

Pool, J. L., & Ransohoff, J. (1949). Autonomic effects on stimulating rostral portion of cingulate gyri in man. Journal of neurophysiology, 12(6), 385–392.

Raut, R. V., Snyder, A. Z., Mitra, A., Yellin, D., Fujii, N., Malach, R., & Raichle, M. E. (2021). Global waves synchronize the brain’s functional systems with fluctuating arousal. Science advances, 7(30), eabf2709.

Raz, S., & Lahad, M. (2022). Physiological indicators of emotional arousal related to ANS activity in response to associative cards for psychotherapeutic PTSD treatment. Frontiers in Psychiatry, 13.

Russo, M. A., Santarelli, D. M., & O’Rourke, D. (2017). The physiological effects of slow breathing in the healthy human. Breathe, 13(4), 298–309.

Saliasi, E., Geerligs, L., Lorist, M. M., & Maurits, N. M. (2014). Neural correlates associated with successful working memory performance in older adults as revealed by spatial ICA. PLoS One, 9(6), e99250.

Salsman, J. M., Butt, Z., Pilkonis, P. A., Cyranowski, J. M., Zill, N., Hendrie, H. C.,… & Cella, D. (2013). Emotion assessment using the NIH Toolbox. Neurology, 80(11 Supplement 3), S76-S86.

Sanbonmatsu, D. M., & Kardes, F. R. (1988). The effects of physiological arousal on information processing and persuasion. Journal of Consumer research, 15(3), 379–385.

Sara, S. J. (2009). The locus coeruleus and noradrenergic modulation of cognition. Nature reviews neuroscience, 10(3), 211–223.

Sara, S. J., & Bouret, S. (2012). Orienting and reorienting: the locus coeruleus mediates cognition through arousal. Neuron, 76(1), 130–141.

Satpute, A. B., Kragel, P. A., Barrett, L. F., Wager, T. D., & Bianciardi, M. (2019). Deconstructing arousal into wakeful, autonomic and affective varieties. Neuroscience letters, 693, 19–28.

Sawai, H., Inou, G., & Koyama, E. (2015, July). Evaluating optimal arousal level during the task based on performance and positive mood: Extracting indices reflecting the relationship among arousal, performance, and mood. In 2015 3rd International Conference on Applied Computing and Information Technology/2nd International Conference on Computational Science and Intelligence (pp. 213-218). IEEE.

Seijdel, N., Schoffelen, J. M., Hagoort, P., & Drijvers, L. (2023). Attention drives visual processing and audiovisual integration during multimodal communication. bioRxiv, 2023–05.

Serrien, D. J., Pogosyan, A. H., & Brown, P. (2004). Influence of working memory on patterns of motor related cortico-cortical coupling. Experimental brain research, 155, 204–210.

Seth, A. K., Barrett, A. B., & Barnett, L. (2015). Granger causality analysis in neuroscience and neuroimaging. Journal of Neuroscience, 35(8), 3293–3297.

Sharot, T., & Phelps, E. A. (2004). How arousal modulates memory: Disentangling the effects of attention and retention. Cognitive, Affective, & Behavioral Neuroscience, 4, 294–306.

Shepherd, L., & Wild, J. (2014). Emotion regulation, physiological arousal and PTSD symptoms in trauma-exposed individuals. Journal of behavior therapy and experimental psychiatry, 45(3), 360–367.

Sonkusare, S., Breakspear, M., & Guo, C. (2019). Naturalistic stimuli in neuroscience: critically acclaimed. Trends in cognitive sciences, 23(8), 699–714.

Spreng, R. N., Shoemaker, L., & Turner, G. R. (2017). Executive functions and neurocognitive aging. In Executive functions in health and disease (pp. 169-196). Academic Press.

Steimer, T. (2022). The biology of fear-and anxiety-related behaviors. Dialogues in clinical neuroscience.

Stockhorst, U., Klosterhalfen, S., Klosterhalfen, W., Winkelmann, M., & Steingrueber, H. J. (1993). Anticipatory nausea in cancer patients receiving chemotherapy: classical conditioning etiology and therapeutical implications. Integrative physiological and behavioral science, 28, 177–181.

Storbeck, J., & Clore, G. L. (2008). Affective arousal as information: How affective arousal influences judgments, learning, and memory. Social and personality psychology compass, 2(5), 1824–1843.

Tracy, J. I., Mohamed, F., Faro, S., Tiver, R., Pinus, A., Bloomer, C.,… & Harvan, J. (2000). The effect of autonomic arousal on attentional focus. Neuroreport, 11(18), 4037–4042.

Vanderwal, T., Eilbott, J., & Castellanos, F. X. (2019). Movies in the magnet: Naturalistic paradigms in developmental functional neuroimaging. Developmental cognitive neuroscience, 36, 100600.

Vergallito, A., Petilli, M. A., Cattaneo, L., & Marelli, M. (2019). Somatic and visceral effects of word valence, arousal and concreteness in a continuum lexical space. Scientific Reports, 9(1), 1–10.

Webb, M, (Director). (2009). (500) Days of Summer [Motion picture]. United States: Fox Searchlight Pictures.

