## Supplementary Materials for "Trait level somatic arousal modulates fMRI neural synchrony to naturalistic stimuli"

**Supplementary Materials A**

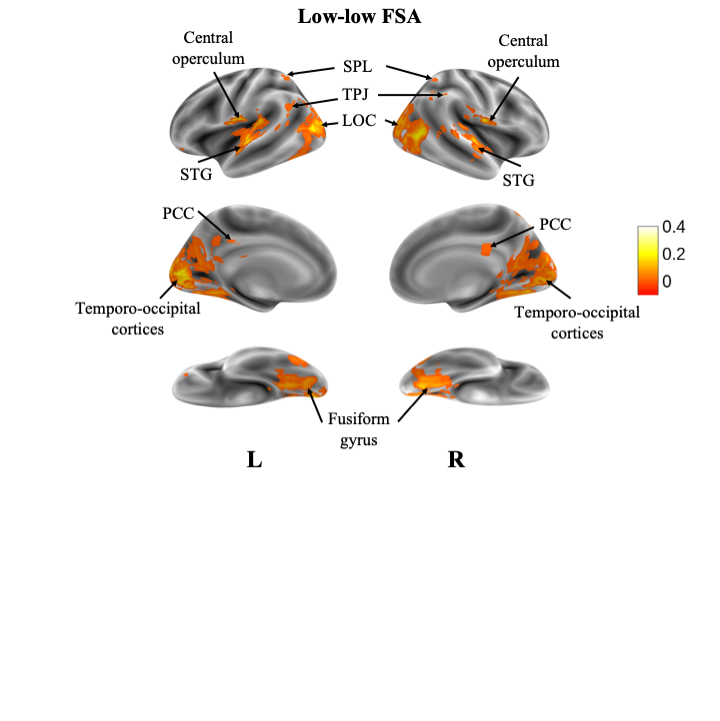

**Figure 1.** Voxels showing significant ISC across the time course of the audiovisual stimulus in participants within the low-low FSA group (*n* = 10). Results are displayed as a voxelwise false-discovery rate (FDR) threshold of *q* = 0.0001

**Table 1**

*Coordinates and cluster sizes of neural synchrony found within the low-low FSA group average contrast. Clusters with five or more voxels shown.*

| **Anatomical Location** | **Hemisphere** | **# of Voxels** | **MAX *r*** | **MAX x** | **MAX y** | **MAX z** |
| --- | --- | --- | --- | --- | --- | --- |
| Intracalcarine Cortex | Left | 502 | 0.242 | -10.5 | -82.5 | 7.5 |
| Lateral Occipital Cortex, inferior division | Right | 298 | 0.199 | 52.5 | -64.5 | 7.5 |
| Lateral Occipital Cortex, inferior division | Left | 142 | 0.241 | -49.5 | -70.5 | 10.5 |
| Occipital Fusiform Gyrus | Right | 135 | 0.147 | 22.5 | -70.5 | -7.5 |
| Temporal Occipital Fusiform Cortex | Left | 100 | 0.126 | -28.5 | -55.5 | -7.5 |
| Planum Temporale | Left | 99 | 0.327 | -61.5 | -16.5 | 7.5 |
| Central Opercular Cortex | Right | 52 | 0.247 | 58.5 | -10.5 | 7.5 |
| Lateral Occipital Cortex, inferior division | Left | 20 | 0.0907 | -49.5 | -67.5 | -10.5 |
| Lateral Occipital Cortex, superior division | Right | 11 | 0.0779 | 13.5 | -58.5 | 64.5 |
| Occipital Pole | Left | 9 | 0.17 | -13.5 | -91.5 | 31.5 |
| Occipital Pole | Right | 8 | 0.131 | 10.5 | -94.5 | 19.5 |
| Superior Temporal Gyrus, anterior division | Left | 7 | 0.182 | -61.5 | -4.5 | -4.5 |
| Lateral Occipital Cortex, superior division | Right | 6 | 0.0569 | 34.5 | -58.5 | 61.5 |
| Lateral Occipital Cortex, superior division | Left | 6 | 0.0631 | -55.5 | -61.5 | 25.5 |
| Lateral Occipital Cortex, superior division | Left | 5 | 0.0794 | -31.5 | -61.5 | 61.5 |
| Lateral Occipital Cortex, superior division | Left | 5 | 0.0611 | -22.5 | -64.5 | 64.5 |

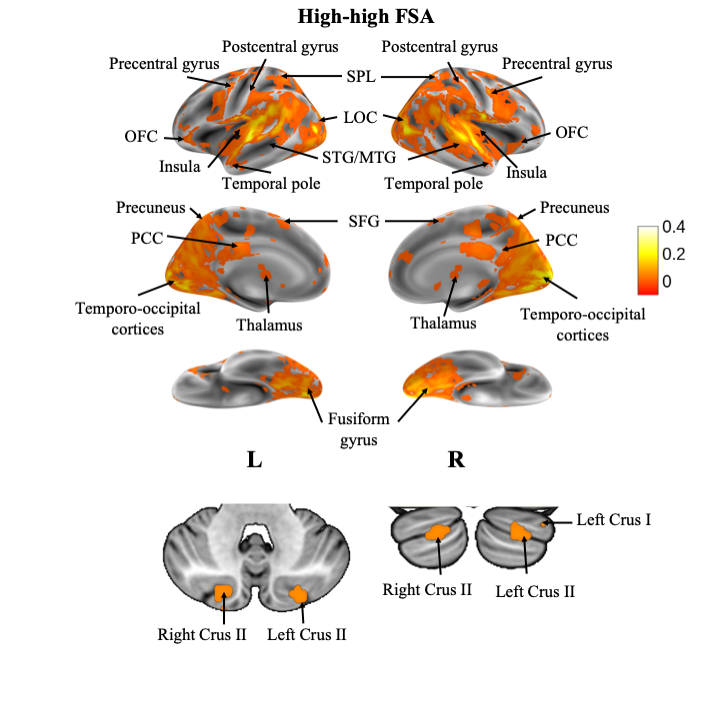

**Figure 2.** Voxels showing significant ISC across the time course of the audiovisual stimulus in participants within the high-high FSA group (*n* = 10). Results are displayed as a voxelwise false-discovery rate (FDR) threshold of *q* = 0.0001

**Table 2**

*Coordinates and cluster sizes of neural synchrony found within the high-high FSA group average contrast. Clusters with five or more voxels shown.*

| Anatomical Location | Hemisphere | # of Voxels | MAX *r* | MAX x | MAX y | MAX z |
| --- | --- | --- | --- | --- | --- | --- |
| Planum Temporale | Right | 5362 | 0.355 | 61.5 | -7.5 | 4.5 |
| Planum Temporale | Left | 774 | 0.361 | -61.5 | -13.5 | 7.5 |
| Precentral Gyrus | Right | 130 | 0.0873 | 49.5 | 10.5 | 34.5 |
| Precentral Gyrus | Right | 87 | 0.0741 | 28.5 | -7.5 | 64.5 |
| Postcentral Gyrus | Left | 53 | 0.0623 | -64.5 | -22.5 | 31.5 |
| Cingulate Gyrus, posterior division | Right | 49 | 0.0485 | 1.5 | -31.5 | 28.5 |
| Precentral Gyrus | Left | 47 | 0.0815 | -40.5 | 1.5 | 34.5 |
| Frontal Pole | Right | 39 | 0.0359 | 28.5 | 40.5 | 43.5 |
| Precentral Gyrus | Left | 37 | 0.062 | -28.5 | -7.5 | 61.5 |
| Supramarginal Gyrus, posterior division | Right | 32 | 0.0438 | 61.5 | -43.5 | 34.5 |
| Cingulate Gyrus, posterior division | Right | 23 | 0.0572 | 13.5 | -22.5 | 40.5 |
| Precentral Gyrus | Right | 22 | 0.0275 | 37.5 | -22.5 | 67.5 |
| Precentral Gyrus | Right | 19 | 0.128 | 58.5 | -1.5 | 46.5 |
| Juxtapositional Lobule Cortex | Left | 19 | 0.0532 | -1.5 | 1.5 | 67.5 |
| Frontal Pole | Right | 18 | 0.047 | 49.5 | 40.5 | 4.5 |
| Frontal Orbital Cortex | Right | 16 | 0.0265 | 31.5 | 25.5 | -4.5 |
| Precentral Gyrus | Left | 16 | 0.123 | -52.5 | -4.5 | 52.5 |
| Precentral Gyrus | Left | 15 | 0.0267 | -43.5 | -13.5 | 58.5 |
| Right Crus II | Right | 14 | 0.044 | 16.5 | -82.5 | -31.5 |
| Superior Frontal Gyrus | Right | 14 | 0.0332 | 28.5 | 22.5 | 58.5 |
| Superior Frontal Gyrus | Left | 13 | 0.0296 | -4.5 | 13.5 | 55.5 |
| Left Crus I | Left | 12 | 0.0493 | -22.5 | -85.5 | -31.5 |
| Caudate | Right | 11 | 0.0211 | 7.5 | 7.5 | 4.5 |
| Frontal Pole | Left | 10 | 0.0353 | -46.5 | 40.5 | 10.5 |
| Caudate | Left | 10 | 0.0171 | -7.5 | 4.5 | 4.5 |
| Superior Frontal Gyrus | Left | 9 | 0.0294 | -25.5 | 19.5 | 55.5 |
| Frontal Pole | Left | 9 | 0.039 | -1.5 | 61.5 | 25.5 |
| Precuneous Cortex | Left | 8 | 0.0708 | -10.5 | -46.5 | 55.5 |
| Precentral Gyrus | Left | 7 | 0.0238 | -7.5 | -28.5 | 79.5 |
| Temporal Fusiform Cortex, posterior division | Right | 7 | 0.0452 | 34.5 | -34.5 | -16.5 |
| Cingulate Gyrus, posterior division | Left | 6 | 0.0318 | -4.5 | -40.5 | 25.5 |
| Insular Cortex | Right | 6 | 0.0329 | 40.5 | -7.5 | -10.5 |
| Precentral Gyrus | Right | 5 | 0.0249 | 10.5 | -31.5 | 73.5 |
| Superior Parietal Lobule | Right | 5 | 0.0736 | 31.5 | -43.5 | 58.5 |
| Right Thalamus | Right | 5 | 0.0202 | 16.5 | -31.5 | -1.5 |
| Middle Frontal Gyrus | Right | 5 | 0.0554 | 52.5 | 25.5 | 25.5 |
| Precentral Gyrus | Left | 5 | 0.0173 | -22.5 | -28.5 | 61.5 |

**Supplementary Materials B**

**Table 3**

*Coordinates and cluster sizes of neural synchrony found within the low-low FSA group compared to the high-high FSA group. Clusters with five or more voxels shown.*

| **Anatomical Location** | **Hemisphere** | **# of Voxels** | **MAX *r*** | **MAX x** | **MAX y** | **MAX z** |
| --- | --- | --- | --- | --- | --- | --- |
| Lateral Occipital Cortex, superior division | Left | 119 | 0.206 | -34.5 | -85.5 | 7.5 |
| Occipital Pole | Left | 45 | 0.212 | -4.5 | -91.5 | 16.5 |
| Lateral Occipital Cortex, inferior division | Right | 40 | 0.198 | 49.5 | -67.5 | -1.5 |
| Lateral Occipital Cortex, inferior division | Right | 37 | 0.195 | 55.5 | -64.5 | 7.5 |
| Central Opercular Cortex | Left | 33 | 0.294 | -61.5 | -16.5 | 10.5 |
| Occipital Pole | Left | 20 | 0.184 | -13.5 | -91.5 | 28.5 |
| Superior Temporal Gyrus, posterior division | Right | 19 | 0.164 | 61.5 | -10.5 | -4.5 |
| Intracalcarine Cortex | Left | 13 | 0.24 | -10.5 | -79.5 | 7.5 |
| Temporal Occipital Fusiform Cortex | Left | 12 | 0.125 | -25.5 | -55.5 | -10.5 |
| Lateral Occipital Cortex, inferior division | Right | 11 | 0.14 | 34.5 | -85.5 | 7.5 |
| Occipital Fusiform Gyrus | Right | 11 | 0.147 | 22.5 | -70.5 | -7.5 |
| Central Opercular Cortex | Left | 10 | 0.203 | -52.5 | -7.5 | 4.5 |
| Lateral Occipital Cortex, superior division | Left | 10 | 0.105 | -13.5 | -82.5 | 43.5 |
| Intracalcarine Cortex | Left | 9 | 0.0979 | -7.5 | -67.5 | 7.5 |
| Superior Parietal Lobule | Left | 8 | 0.073 | -34.5 | -52.5 | 64.5 |
| Lateral Occipital Cortex, inferior division | Left | 8 | 0.106 | -40.5 | -67.5 | 13.5 |
| Lateral Occipital Cortex, superior division | Right | 7 | 0.115 | 49.5 | -73.5 | 19.5 |
| Lateral Occipital Cortex, superior division | Right | 7 | 0.0752 | 13.5 | -58.5 | 61.5 |
| Supramarginal Gyrus, posterior division | Right | 7 | 0.0778 | 64.5 | -37.5 | 22.5 |
| Superior Temporal Gyrus, anterior division | Left | 6 | 0.14 | -58.5 | -4.5 | -7.5 |
| Superior Temporal Gyrus, posterior division | Left | 6 | 0.121 | -61.5 | -19.5 | -1.5 |
| Inferior Temporal Gyrus, temporooccipital part | Left | 6 | 0.0644 | -43.5 | -58.5 | -7.5 |
| Angular Gyrus | Left | 6 | 0.0577 | -55.5 | -55.5 | 25.5 |
| Occipital Pole | Right | 5 | 0.156 | 22.5 | -91.5 | 22.5 |
| Occipital Fusiform Gyrus | Left | 5 | 0.114 | -25.5 | -73.5 | -4.5 |
| Parahippocampal Gyrus, posterior division | Left | 5 | 0.0409 | -13.5 | -40.5 | -7.5 |

**Table 4**

*Coordinates and cluster sizes of neural synchrony found within the high-high FSA group compared to the low-low FSA group. Clusters with five or more voxels shown.*

| **Anatomical Location** | **Hemisphere** | **# of Voxels** | **MAX r** | **MAX x** | **MAX y** | **MAX z** |
| --- | --- | --- | --- | --- | --- | --- |
| Occipital Pole | Right | 1280 | 0.292 | 10.5 | -91.5 | 1.5 |
| Planum Temporale | Right | 559 | 0.355 | 61.5 | -7.5 | 4.5 |
| Planum Temporale | Left | 467 | 0.343 | -55.5 | -19.5 | 7.5 |
| Superior Temporal Gyrus, posterior division | Left | 232 | 0.224 | -67.5 | -13.5 | 4.5 |
| Lateral Occipital Cortex, superior division | Left | 226 | 0.163 | -31.5 | -88.5 | 16.5 |
| Precentral Gyrus | Right | 183 | 0.0741 | 28.5 | -7.5 | 64.5 |
| Supramarginal Gyrus, anterior division | Right | 153 | 0.0822 | 49.5 | -34.5 | 55.5 |
| Temporal Occipital Fusiform Cortex | Right | 110 | 0.143 | 31.5 | -46.5 | -7.5 |
| Lateral Occipital Cortex, superior division | Right | 107 | 0.12 | 37.5 | -82.5 | 25.5 |
| Precentral Gyrus | Right | 86 | 0.0873 | 49.5 | 10.5 | 34.5 |
| Lingual Gyrus | Left | 69 | 0.195 | -10.5 | -79.5 | -7.5 |
| Temporal Occipital Fusiform Cortex | Left | 49 | 0.117 | -28.5 | -55.5 | -10.5 |
| Precentral Gyrus | Left | 32 | 0.123 | -52.5 | -4.5 | 52.5 |
| Superior Parietal Lobule | Left | 32 | 0.0749 | -37.5 | -52.5 | 58.5 |
| Superior Frontal Gyrus | Left | 31 | 0.0451 | -4.5 | 16.5 | 64.5 |
| Precentral Gyrus | Right | 29 | 0.128 | 58.5 | -1.5 | 46.5 |
| Precentral Gyrus | Left | 27 | 0.062 | -28.5 | -7.5 | 61.5 |
| Lateral Occipital Cortex, inferior division | Left | 27 | 0.184 | -49.5 | -76.5 | -1.5 |
| Lateral Occipital Cortex, inferior division | Right | 25 | 0.265 | 52.5 | -70.5 | 4.5 |
| Precentral Gyrus | Left | 24 | 0.0931 | -43.5 | 1.5 | 34.5 |
| Precentral Gyrus | Left | 22 | 0.027 | -46.5 | -10.5 | 58.5 |
| Frontal Pole | Right | 19 | 0.047 | 49.5 | 40.5 | 4.5 |
| Lateral Occipital Cortex, superior division | Left | 19 | 0.124 | -28.5 | -88.5 | 25.5 |
| Frontal Pole | Left | 18 | 0.039 | -1.5 | 61.5 | 25.5 |
| Cingulate Gyrus, posterior division | Right | 17 | 0.0572 | 13.5 | -22.5 | 40.5 |
| Superior Parietal Lobule | Right | 15 | 0.0736 | 31.5 | -43.5 | 58.5 |
| Angular Gyrus | Left | 14 | 0.0718 | -46.5 | -58.5 | 25.5 |
| Frontal Pole | Right | 14 | 0.0359 | 31.5 | 55.5 | 28.5 |
| Frontal Pole | Right | 14 | 0.0344 | 28.5 | 46.5 | 40.5 |
| Temporal Pole | Right | 12 | 0.0418 | 49.5 | 25.5 | -22.5 |
| Middle Temporal Gyrus, anterior division | Left | 10 | 0.0214 | -55.5 | -1.5 | -34.5 |
| Left Crus II | Left | 10 | 0.0508 | -22.5 | -88.5 | -31.5 |
| Lateral Occipital Cortex, superior division | Right | 10 | 0.0399 | 52.5 | -64.5 | 31.5 |
| Right Crus II | Right | 10 | 0.044 | 16.5 | -82.5 | -31.5 |
| Postcentral Gyrus | Right | 10 | 0.026 | 37.5 | -25.5 | 61.5 |
| Precentral Gyrus | Left | 9 | 0.0431 | -13.5 | -19.5 | 40.5 |
| Superior Frontal Gyrus | Left | 9 | 0.0294 | -25.5 | 19.5 | 55.5 |
| Precuneous Cortex | Right | 9 | 0.07 | 10.5 | -46.5 | 58.5 |
| Frontal Pole | Left | 9 | 0.0331 | -25.5 | 40.5 | 43.5 |
| Insular Cortex | Right | 9 | 0.0329 | 40.5 | -7.5 | -10.5 |
| Superior Frontal Gyrus | Right | 8 | 0.0332 | 28.5 | 22.5 | 58.5 |
| Planum Polare | Left | 8 | 0.0251 | -46.5 | 1.5 | -13.5 |
| Precuneous Cortex | Left | 7 | 0.112 | -19.5 | -61.5 | 19.5 |
| Left VI | Left | 7 | 0.0144 | -34.5 | -43.5 | -40.5 |
| Frontal Orbital Cortex | Right | 7 | 0.0265 | 31.5 | 25.5 | -4.5 |
| Left I-IV | Left | 6 | 0.0135 | -1.5 | -40.5 | 1.5 |
| Middle Frontal Gyrus | Left | 6 | 0.0284 | -28.5 | 25.5 | 49.5 |
| Precentral Gyrus | Left | 5 | 0.0238 | -7.5 | -28.5 | 79.5 |
| Cuneal Cortex | Left | 5 | 0.0939 | -4.5 | -88.5 | 31.5 |
| Postcentral Gyrus | Right | 5 | 0.0213 | 19.5 | -28.5 | 76.5 |
| Temporal Occipital Fusiform Cortex | Left | 5 | 0.06 | -46.5 | -58.5 | -19.5 |
| Frontal Pole | Left | 5 | 0.0219 | -25.5 | 52.5 | 34.5 |
| Precuneous Cortex | Left | 5 | 0.0819 | -7.5 | -73.5 | 37.5 |

**Supplementary Materials C**

**Table 5**

*Coordinates and cluster sizes of neural synchrony found within the low-low FSA group compared to the low-high FSA group. Clusters with five or more voxels shown.*

| **Anatomical Location** | **Hemisphere** | **# of Voxels** | **MAX *r*** | **MAX x** | **MAX y** | **MAX z** |
| --- | --- | --- | --- | --- | --- | --- |
| Planum Temporale | Left | 1 | 0.176 | -64.5 | -19.5 | 7.5 |
| Occipital Pole | Left | 1 | 0.0745 | -13.5 | -94.5 | 34.5 |

**Table 6**

*Coordinates and cluster sizes of neural synchrony found within the high-high FSA group compared to the low-high FSA group. Clusters with five or more voxels shown.*

| **Anatomical Location** | **Hemisphere** | **# of Voxels** | **MAX *r*** | **MAX x** | **MAX y** | **MAX z** |
| --- | --- | --- | --- | --- | --- | --- |
| Occipital Pole | Right | 287 | 0.292 | 10.5 | -91.5 | 1.5 |
| Planum Temporale | Right | 235 | 0.355 | 61.5 | -7.5 | 4.5 |
| Heschl's Gyrus | Left | 105 | 0.315 | -52.5 | -22.5 | 7.5 |
| Lingual Gyrus | Right | 89 | 0.178 | 10.5 | -76.5 | -10.5 |
| Precentral Gyrus | Right | 74 | 0.0741 | 28.5 | -7.5 | 64.5 |
| Supramarginal Gyrus, anterior division | Right | 71 | 0.0822 | 49.5 | -34.5 | 55.5 |
| Supramarginal Gyrus, posterior division | Left | 68 | 0.187 | -67.5 | -43.5 | 7.5 |
| Superior Temporal Gyrus, anterior division | Left | 63 | 0.177 | -61.5 | 1.5 | -1.5 |
| Lateral Occipital Cortex, superior division | Right | 59 | 0.114 | 7.5 | -61.5 | 67.5 |
| Frontal Operculum Cortex | Left | 53 | 0.0815 | -46.5 | 28.5 | -4.5 |
| Heschl's Gyrus | Right | 46 | 0.193 | 49.5 | -16.5 | 10.5 |
| Precentral Gyrus | Right | 44 | 0.0873 | 49.5 | 10.5 | 34.5 |
| Lateral Occipital Cortex, inferior division | Left | 39 | 0.151 | -34.5 | -91.5 | -1.5 |
| Precentral Gyrus | Right | 37 | 0.128 | 58.5 | -1.5 | 46.5 |
| Angular Gyrus | Right | 31 | 0.152 | 58.5 | -46.5 | 19.5 |
| Postcentral Gyrus | Left | 30 | 0.0623 | -64.5 | -22.5 | 31.5 |
| Precuneous Cortex | Right | 30 | 0.0968 | 1.5 | -61.5 | 37.5 |
| Lateral Occipital Cortex, superior division | Left | 28 | 0.103 | -19.5 | -73.5 | 55.5 |
| Lingual Gyrus | Left | 22 | 0.193 | -7.5 | -76.5 | -1.5 |
| Occipital Pole | Left | 21 | 0.162 | -13.5 | -100 | 1.5 |
| Juxtapositional Lobule Cortex | Left | 19 | 0.0532 | -1.5 | 1.5 | 67.5 |
| Precuneous Cortex | Right | 17 | 0.153 | 1.5 | -58.5 | 55.5 |
| Temporal Occipital Fusiform Cortex | Right | 15 | 0.0865 | 31.5 | -46.5 | -19.5 |
| Superior Parietal Lobule | Left | 15 | 0.0492 | -43.5 | -43.5 | 52.5 |
| Superior Frontal Gyrus | Left | 14 | 0.0451 | -4.5 | 16.5 | 64.5 |
| Lateral Occipital Cortex, superior division | Left | 14 | 0.0912 | -22.5 | -67.5 | 49.5 |
| Precentral Gyrus | Left | 14 | 0.123 | -52.5 | -4.5 | 52.5 |
| Lateral Occipital Cortex, inferior division | Left | 13 | 0.18 | -46.5 | -76.5 | -1.5 |
| Precentral Gyrus | Left | 12 | 0.0931 | -43.5 | 1.5 | 34.5 |
| Frontal Pole | Right | 12 | 0.0344 | 28.5 | 46.5 | 40.5 |
| Temporal Occipital Fusiform Cortex | Left | 11 | 0.0803 | -31.5 | -58.5 | -10.5 |
| Superior Parietal Lobule | Left | 10 | 0.102 | -16.5 | -58.5 | 67.5 |
| Frontal Pole | Right | 10 | 0.047 | 49.5 | 40.5 | 4.5 |
| Superior Temporal Gyrus, posterior division | Left | 9 | 0.203 | -67.5 | -25.5 | 1.5 |
| Precentral Gyrus | Left | 9 | 0.0616 | -31.5 | -7.5 | 61.5 |
| Occipital Pole | Left | 9 | 0.147 | -7.5 | -91.5 | 10.5 |
| Lateral Occipital Cortex, superior division | Right | 9 | 0.0869 | 31.5 | -67.5 | 49.5 |
| Frontal Pole | Left | 9 | 0.039 | -1.5 | 61.5 | 25.5 |
| Precuneous Cortex | Right | 9 | 0.104 | 10.5 | -67.5 | 37.5 |
| Occipital Pole | Right | 8 | 0.123 | 37.5 | -91.5 | -7.5 |
| Temporal Occipital Fusiform Cortex | Right | 8 | 0.143 | 31.5 | -46.5 | -7.5 |
| Occipital Fusiform Gyrus | Right | 8 | 0.101 | 25.5 | -79.5 | -4.5 |
| Lateral Occipital Cortex, inferior division | Right | 8 | 0.265 | 52.5 | -70.5 | 4.5 |
| Temporal Occipital Fusiform Cortex | Right | 7 | 0.0858 | 43.5 | -49.5 | -22.5 |
| Lateral Occipital Cortex, superior division | Right | 7 | 0.12 | 37.5 | -82.5 | 25.5 |
| Frontal Pole | Right | 6 | 0.0359 | 31.5 | 55.5 | 28.5 |
| Superior Temporal Gyrus, posterior division | Right | 6 | 0.139 | 67.5 | -25.5 | 16.5 |
| Occipital Fusiform Gyrus | Left | 6 | 0.0979 | -31.5 | -79.5 | -16.5 |
| Precuneous Cortex | Left | 5 | 0.0725 | -13.5 | -52.5 | 4.5 |
| Superior Parietal Lobule | Right | 5 | 0.0736 | 31.5 | -43.5 | 58.5 |
| Cingulate Gyrus, posterior division | Right | 5 | 0.0572 | 13.5 | -22.5 | 40.5 |
| Frontal Orbital Cortex | Right | 5 | 0.0265 | 31.5 | 25.5 | -4.5 |
